# Genome organization in double-stranded DNA viruses observed by cryoET

**DOI:** 10.1101/2023.12.15.571939

**Authors:** Muyuan Chen, Bibekananda Sahoo, Zongjun Mou, Xiyong Song, Tiffany Tsai, Xinghong Dai

**Affiliations:** Division of CryoEM and Bioimaging, SSRL, SLAC National Accelerator Laboratory, Stanford University, Menlo Park, CA 94025, USA; Department of Physiology and Biophysics, Case Western Reserve University, Cleveland, OH 44106, USA

## Abstract

Double-stranded DNA (dsDNA) viruses package their genetic material into protein cages with diameters usually a few hundred times smaller than the length of their genome. Compressing the relatively stiff and highly negatively charged dsDNA into a small volume is energetically costly and mechanistically enigmatic. Multiple models of dsDNA packaging have been proposed based on various experimental evidence and simulation methods, but direct observation of any viral genome organization is lacking. Here, using cryoET and an improved data processing scheme that utilizes information from the encaging protein shell, we present 3D views of dsDNA genome inside individual viral particles at resolution that densities of neighboring DNA duplexes are readily separable. These cryoET observations reveal a “rod-and-coil” fold of the dsDNA that is conserved among herpes simplex virus type 1 (HSV-1) with a spherical capsid, bacteriophage T4 with a prolate capsid, and bacteriophage T7 with a proteinaceous core inside the capsid. Finally, inspired by the genome arrangement in partially packaged T4 particles, we propose a mechanism for the genome packaging process in dsDNA viruses.

## Main

Herpesviruses and tailed bacteriophages package their genetic material–a linear, double-stranded DNA (dsDNA)–into a preformed empty capsid (procapsid or prohead) that is only a few hundredths in dimension compared to the length of the dsDNA being packaged. Compressing the relatively stiff and highly negatively charged dsDNA to near-crystalline density inside the capsid results in an internal pressure of ∼20 atmospheres. A powerful, multi-subunit machinery called terminase is responsible for pumping the concatemeric viral DNA—a product of the viral replication—into the capsid through a portal embedded in the capsid shell. The portal is a dodecameric ring with a central channel just big enough for a *linear* dsDNA to thread through^1^. The portal is also responsible for sensing the “headfull” signal, which triggers the terminase to cut the concatemeric DNA, resulting in roughly one unit of the viral genome being packaged. The last bit of the packaged DNA is retained in the channel of the portal, poised for leading the way out when the viral particle infects a new host. This “last in, first out” manner of linear ejection through the portal requires the unwinding of the packaged DNA within the confined space of the capsid lumen. It is enigmatic how the DNA is organized inside the capsid to accommodate dense packaging and fast, tangle-free unwinding at the same time.

The enquiry of dsDNA genome organization in phage heads has been going on for more than 60 years, yet with no concrete answer. As early as 1961, X-ray fiber diffraction of oriented T2 phages had revealed the hexagonal packing of B-form DNA in “crystalline domains” each with 25-30 parallel DNA segments^2^. Negative-staining electron microscopy (EM) of partially ruptured phage heads revealed circular outlines, in some cases concentric circles, of the DNA strands, leading to the proposal of a “ball of yarn” model, and a “coaxial spool” model^3^ that has been most widely accepted^4–6^. Recently, the “coaxial spool” model gained more popularity as cryogenic electron microscopy (cryoEM) and 3D reconstruction of various phage heads and herpesvirus capsids showed concentric layers or even strands of DNA densities in the outermost layers of the spool^7–11^. However, while clear DNA densities could be observed near the inner surface of the capsid, the cryoEM density maps appeared completely disordered at the center of the virus, leading to an incomplete picture of DNA packaging in these cases. In the meantime, the viral DNA packaging problem has also been tackled by molecular dynamics (MD) simulation studies over the decades^12–14^. While the actual system of dsDNA packaging is too large in both size and time scale even for modern MD simulation techniques, simulation results on smaller scales still serve as validations for different packaging models, and bring useful insights in interpreting experimental data.

It had been recognized before that cryogenic electron tomography (cryoET) might be able to offer a definitive solution to this dsDNA packaging enigma, although there were “major, possibly unattainable, challenges” in reaching the required resolution^11^. Recent advances in cryoET computational methods, particularly subtomogram averaging in combination with subtilt refinement, have allowed atomic structure determination of homogeneous components from tilt series of a pleomorphic sample^15–17^. Importantly, this integrative procedure not only pushes averaged structures toward higher resolution, but also allows better visualization of individual particles, including their heterogeneous features. Each dsDNA virus particle is a combination of homogeneous component—a well-defined icosahedral capsid—with heterogeneous component—a packaged genome that is pleomorphic from particle to particle. In this work, we imaged dsDNA viruses with cryoET, and performed subtomogram averaging and subtilt refinement with their capsids. This procedure led to the resolving of separated DNA duplexes in the polished subtomograms, enabling direct observation of the genome organization in dsDNA viruses for the first time.

First, we imaged the virion of human herpes simplex virus type 1 (HSV-1) with cryoET. From the reconstructed tomograms, subtomograms of HSV-1 capsids were segmented and subjected to iterative subtomogram and subtilt refinement (Fig. 1 a-b). Then, the rotation and translation parameters of each subtilt derived from this refinement were used to reconstruct the “polished subtomogram” for each particle. The polished subtomograms had a much-improved signal to noise ratio compared to the raw subtomograms, with DNA duplex densities becoming discernible in the XY planes as threads, dots, or short rods, corresponding to the cross section of a DNA duplex that goes in plane, normal to the plane, or in-between, respectively (Fig. 2, Supplementary video 1-3). Due to the much more limited resolution of tomographic reconstruction in the Z-direction, we were not able to directly trace the DNA duplexes in 3D and build models for individual particles. However, from the planar views of the polished HSV-1 particles, it could be derived that the DNA duplex was organized in a rod-and-coil fold, where a central rod was wrapped around by multi-layer, right-handed coils (Fig. 1b). In some particles where the longitudinal axis of the rod happened to lie in the XY plane, the rod was clearly seen as a bundle of threads traversing the capsid lumen near the center, while the coils were hexagonal arrays of dots on both sides of the rod (Fig. 1c). In average, there were about eight layers of DNA coil surrounding the rod. In some other particles where the rod was nearly perpendicular to the XY plane, the centrally sliced view became an array of dots surrounded by circles (Fig. 1d). Near the two poles of the rod, where the strands in the rod were transitioning to the coils, two vortex patterns were formed (Fig. 1e). Remarkably, the longitudinal axis of the DNA rod was slightly off the central axis of the capsid. This asymmetry in positioning gave more room on one side of the central rod compared to the other, accommodating the two vortices. Surprisingly, unlike what has been generally assumed in the coaxial spool model that the spool axis more or less aligns with the portal axis, the relative orientation between the central rod and the portal axis in HSV-1 seems to be highly variable among the observed particles (Fig. S4). This may explain why the rod-and-coil organization of the dsDNA genome was never observed in previous CryoEM reconstructions of herpesvirus capsids obtained by single particle analysis.

**Fig. 1.**
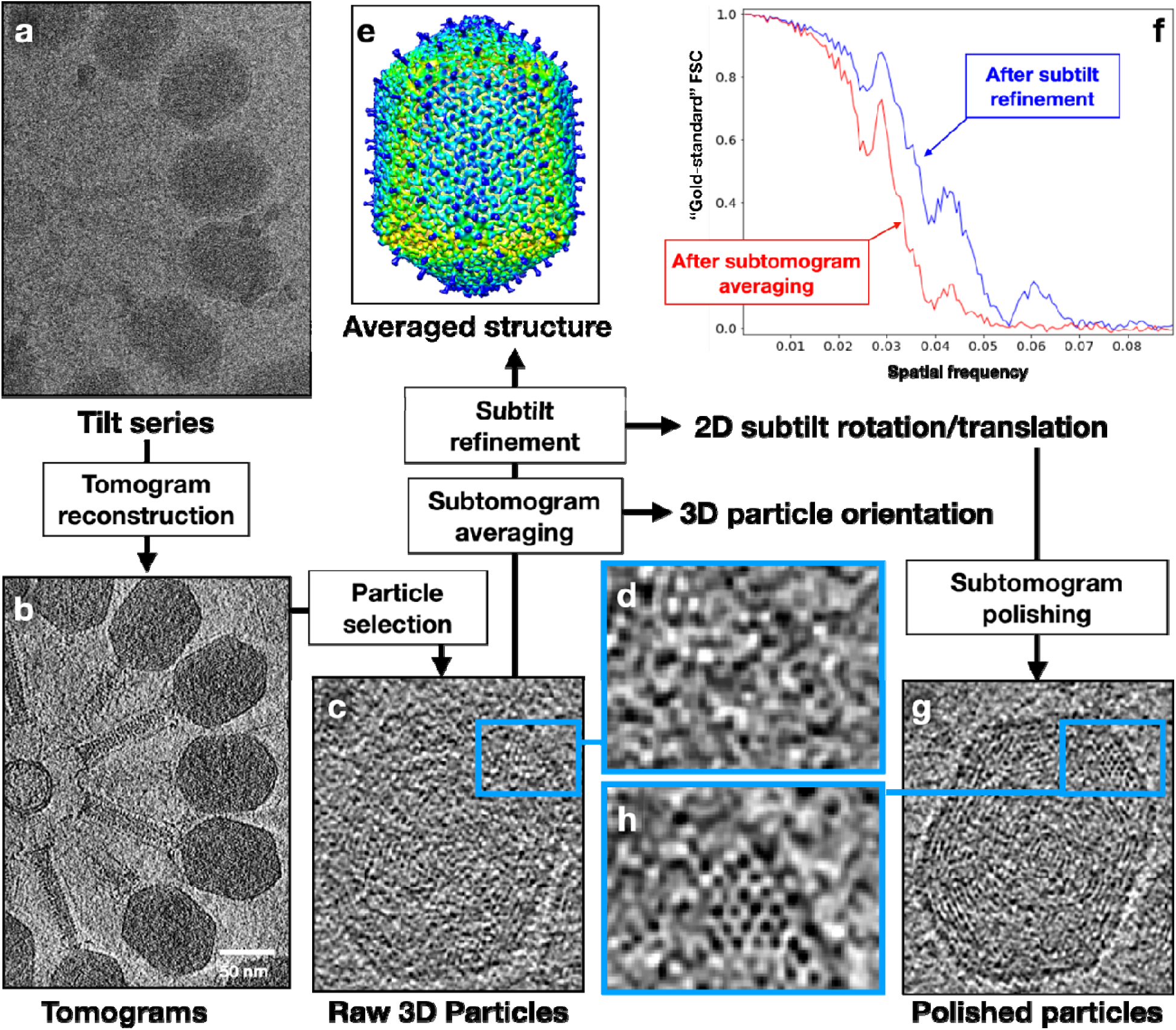
Visualizing DNA packaging inside virus capsids using cryoET and subtomogram polishing. (a) Raw electron micrograph of T4 bacteriophage. (b) Central slice of a reconstructed tomogram from the tilt series. (c) Slice view of one particle from the tomogram. (d) Zoomed in view of the DNA packaging inside capsid from c. (e) Averaged 3D structure of the T4 capsid from 184 particles, after both subtomogram and subtilt refinement, colored by distance from the center of volume. (f) Plot of “gold-standard” FSC showing the improvement of resolution after subtilt refinement. (g) Slice view of the same particle in c, after subtomogram polish using the subtilt refinement result of the capsid. (h) Zoomed in view of the DNA packaging inside capsid from g.

**Fig. 2.**
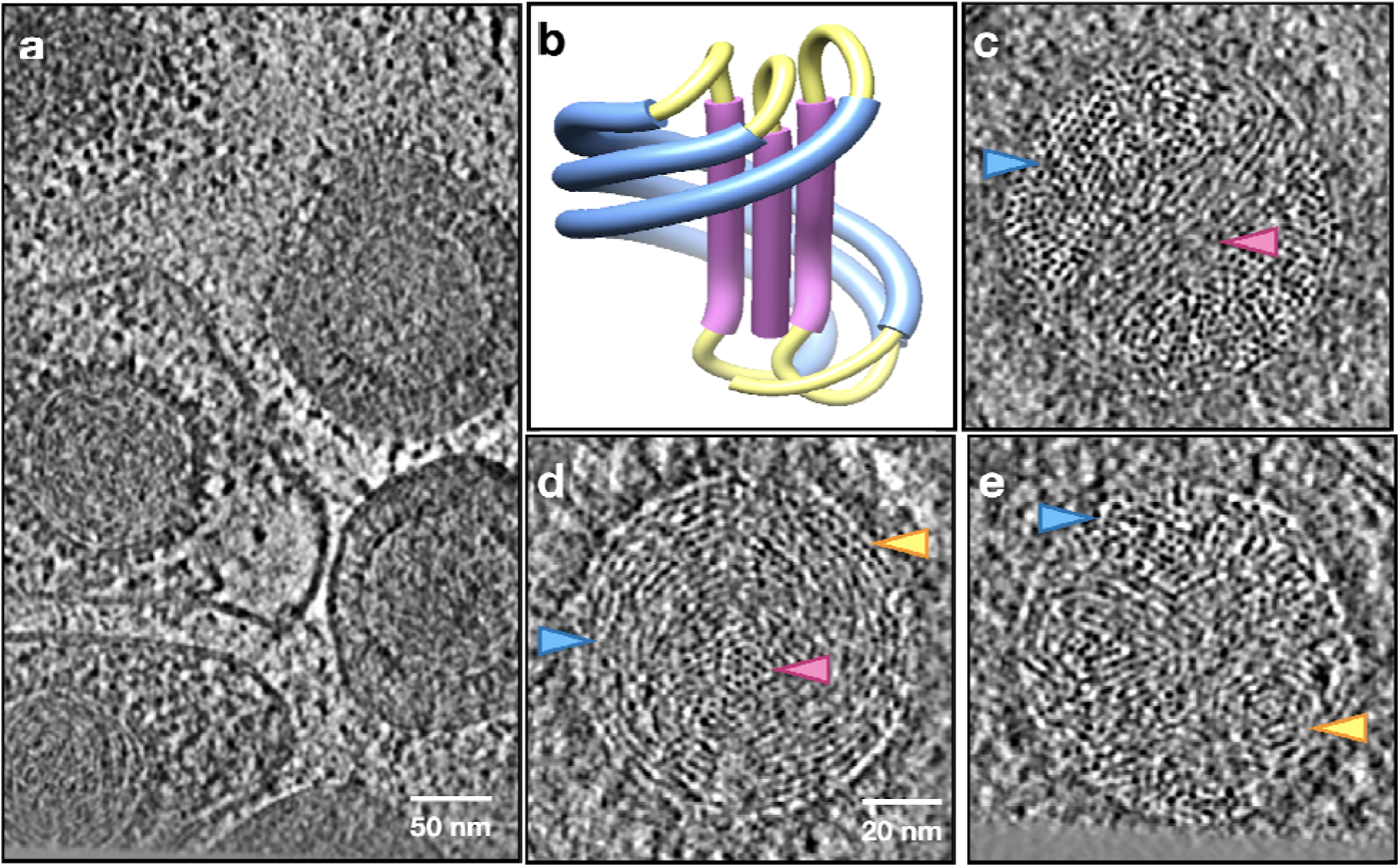
Visualizing DNA packaging in HSV-1 subtomograms. (a) Slice view of a tomogram of purified HSV-1. (b) Diagram of the rod-and-coil model. (c) Slice view of a polished HSV-1 particle whose central DNA rod is on the imaging plane. Arrows: pink - central DNA rod; blue - coil. (d) Slice view of a polished HSV particle whose central DNA rod is perpendicular to the imaging plane. Pink - central rod; blue - coil; yellow - seam line. (e) Slice view of a polished HSV particle, showing the vortex connecting the coil to the central DNA rod. Blue - coil; yellow - vortex.

Locally, the DNA duplexes form tightly packed liquid crystalline, with their cross sections conforming to a hexagonal lattice (Fig. S1). The average center-to-center distance of the nearest DNA duplexes was measured to be 31 Å. This number seems to be contradicting, but is actually consistent with, the 26 Å inter-layer spacing measured previously from single particle cryoEM of HSV-1 capsid^7,18^. In the latter case, the staggered DNA duplexes are rotationally averaged and smeared out as density shells with a shortened inter-layer distance, as illustrated with a simple geometric calculation (Fig. S2).

Given the local hexagonal pattern and 31-Å spacing, we were able to estimate several other important parameters required for modeling the DNA organization. First, we estimated that there were about 88 DNA duplexes in the central rod by measuring the cross-section area of the rod. The ratio of DNA length between the coil and the rod could then be calculated from the volume of the rod and the capsid lumen (Fig S3). This number roughly corresponds to the ratio one would expect when the DNA coil makes two full turns (720 degrees) around the central rod in a simplified model we used for numerical simulation (Fig. S4).

Using these parameters, we built a conceptual model of genome organization inside the HSV-1 capsid (Fig. 3a, Supplementary video 11). Although direct comparison between the model and the subtomograms is challenging due to the limited resolution along the Z direction, thin slices in the XY-plane of the model matched well with those of the subtomograms at different positions throughout the Z-axis (Fig. 2b-c), serving as a strong validation of our model. Notably, in a best match between the model and the subtomograms, the DNA in the coil does not have a uniform rise along the rod throughout its path. For most parts of the coil, the DNA strand only rises slightly, but the rise increases drastically for a small segment, forming the pattern of a seam line. This pattern can be best visualized in particles whose central rod is parallel to the incoming electron beam. In the cross section of those particles, most coil DNA appears as curved threads on the imaging plane, but there is always a small area of the coil where the DNA appears as dots (green in Fig. 2d), suggesting that they are running nearly normal to the plane like those in the central rod. When viewed from an orthogonal direction, the seam line, along with DNA at the outer parts of the central rod, forms the vortexes that bridge the central rod to the coil (Fig. 2e).

**Fig. 3.**
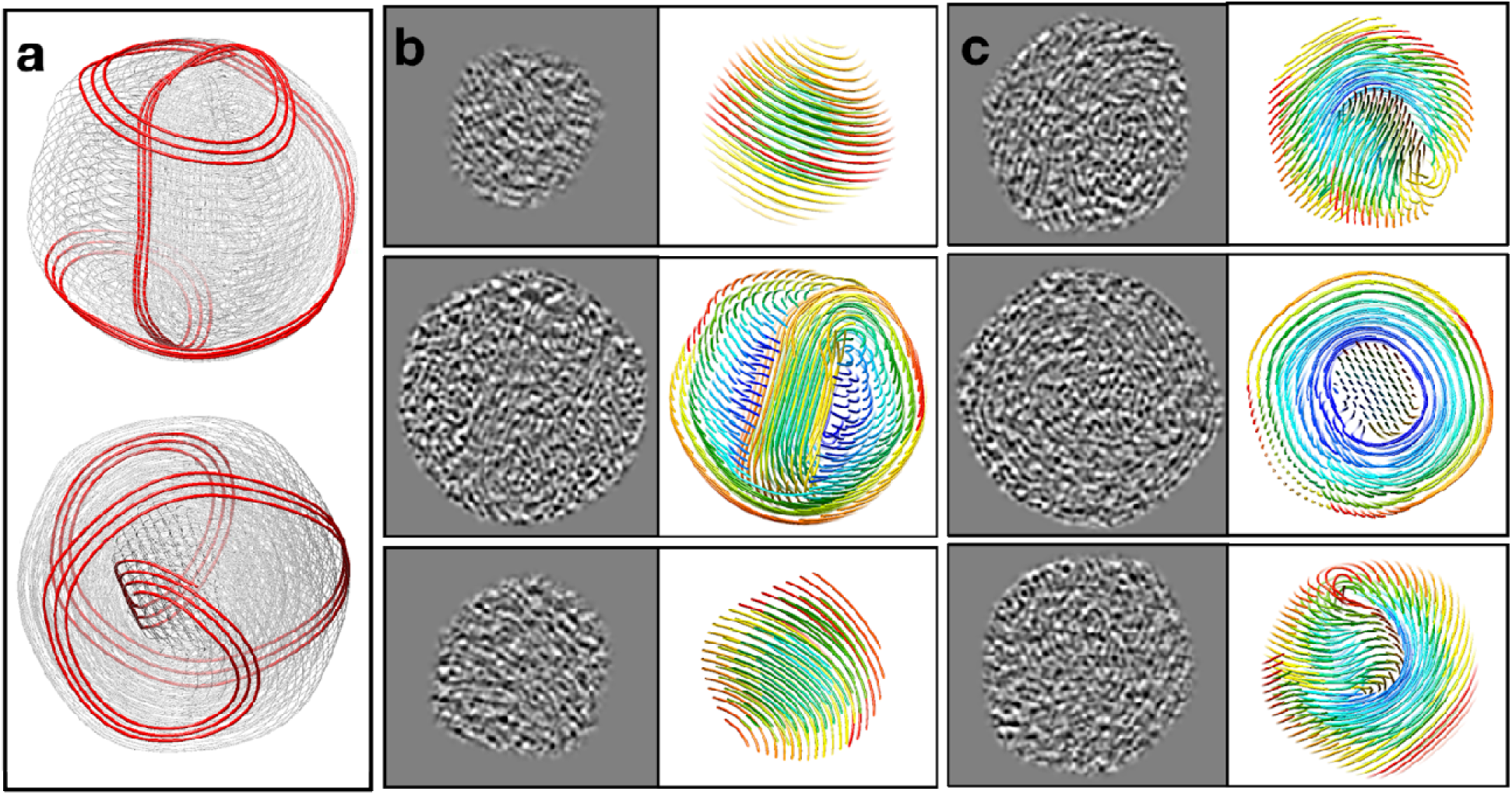
Model of DNA organization and packaging mechanism. (a) Conceptual model of genome organization in HSV-1. (b-c) Slice-by-slice comparison between the cross-section of the model in a, and the slice view of DNA inside polished HSV-1 subtomograms.

It is worth noting that the conceptual model does not fit exactly into every particle, due to the variability of DNA packaging in different particles. For example, as the orientation of the rod and vortex with respect to the capsid shell varies, the DNA in the coil would be shaped by the inner surface of the capsid in different ways, causing differences in DNA traces. In addition, the tilt angle between the equator of the coil and the axis of the rod can also be different among the particles, resulting in differences in the overall fold of DNA.

To generalize the DNA organization model, we imaged T4 bacteriophage, another well-studied dsDNA virus with prolate icosahedral capsid. Polished T4 subtomograms were generated in the same way as those of the HSV-1. The T4 particles had a strong preferred orientation with the long axis of the capsid and the tail lying in the imaging plane. Nonetheless, it was still appreciable from the limited number of views that the T4 DNA is packaged in a similar rod-and-coil fold as that in HSV-1. Signature features, including the central rod and the coil around, as well as the seam line along the coil, were clearly visible in the particles (Fig. 1g, Supplementary video 4-6). The DNA rod sits slightly off the center of the capsid, roughly along the short axis of the capsid and perpendicular to the long axis (Fig. S4). The local packaging of DNA is tighter in T4 compared to that in HSV-1, with an average nearest neighbor distance of 26Å.

Among the most studied dsDNA viruses, bacteriophage T7 has a unique feature that, within the capsid, a prominent, proteinaceous core is attached to the portal and surrounded by the dsDNA genome. During the T7 infection, these internal core proteins are translocated and reassembled into a tube within the periplasm of the host bacteria to extend the tail channel and conduct the genome^19–22^. To see the influence of an internal core on dsDNA genome organization, we imaged T7 phages and performed the same analysis as HSV-1 and T4. In the polished T7 particles, most of the DNA forms concentric coils surrounding the protein core (Fig. 4, Supplementary video 9-10). While the protein core only traverses about half of the capsid lumen, the concentric DNA coils extend beyond the length of the protein core, reaching the pentamer at the other end of the capsid. Overall, the T7 genome organization is the closest to the classic concentric spooling model. Nonetheless, in the space above the protein core and at the center of the DNA coil, a local rod-and-coil pattern can still be observed in some of the particles, when the central rod aligns with the Z direction. The nearest neighbor distance between adjacent DNA duplexes in the concentric coils of T7 was measured to be around 24Å, even smaller than that in HSV-1 and T4.

**Fig. 4.**
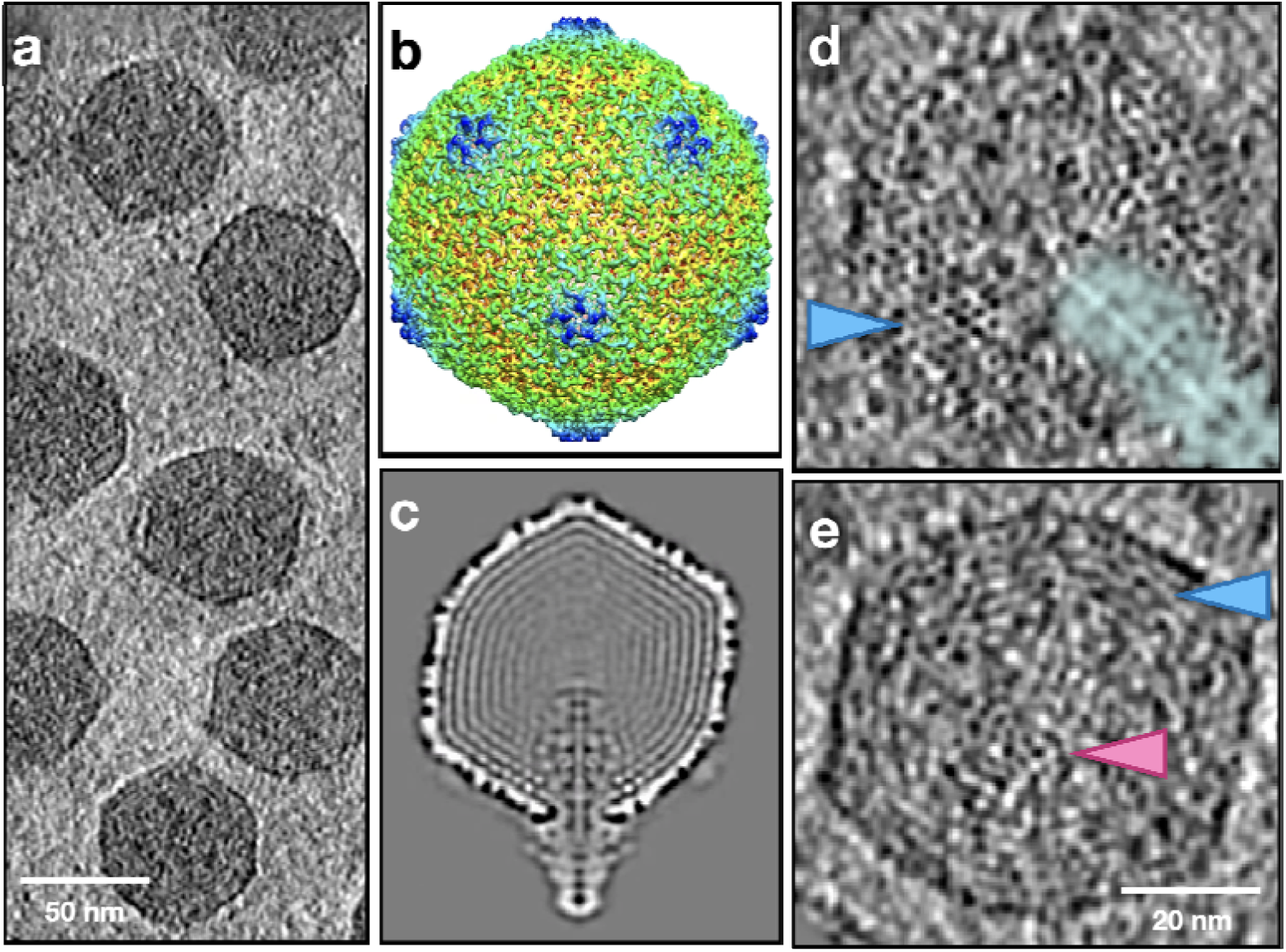
Genome packaging in T7. (a) Slice view of a tomogram of T7. (b) Averaged structure of T7 after subtilt refinement with icosahedral symmetry. (c) Central slice view of the averaged T7 structure without symmetry, showing the protein core inside the capsid. (d) Slice view of a polished T7 particle whose portal axis is on the imaging plane. The density of the protein core is highlighted by the cyan shadow. (e) Slice view of a polished T7 particle whose portal axis is perpendicular to the imaging plane. Arrows: pink - central DNA rod; blue - coil.

In sum, the observations we made from the cryoET datasets of HSV-1 and T4 indicated that the rod-and-coil fold is a common form of genome organization in dsDNA viruses. While most of the DNA is winded around the protein core in T7, the rod-and-coil fold can still form locally at the space above the protein core inside the capsid, suggesting that this fold is a preferred state for DNA to pack in a compact space. As dsDNA is a rigid polymer with a relatively long persistence length (30-100nm) ^23–26^, any segment of the packaged genome would prefer to stay as extended as possible, and sharp bending should be strictly avoided. On the other hand, dsDNA is highly negatively charged; electrostatic repulsion between nearby segments would keep them well separated, or evenly spread out to occupy every bit of space in the capsid lumen when genome packaging is near the end. A prominent example demonstrating the latter scenario can be found in previous CryoEM reconstruction of human herpesvirus 5 (aka human cytomegalovirus, HCMV), where the outermost layer of the packaged genome was even squeezed into the channel of the capsomers^27^. Last but not least, an even distribution of the dsDNA across the entire space of the capsid lumen is entropically favored. Intuitively, compared to many other proposed models, the rod-and-coil model better fits to the above principles *simultaneously*. For example, in a coaxial spooling model, to fill up the space near the axis of the spool, the dsDNA has to be sharply bent; while in the rod-and-coil model, the entire space is evenly occupied, and most dsDNA segments are well-extended with the most significant bending occurs in the two vertex regions.

As the same principles should also apply *during* genome packaging, it is now possible to speculate the intermediate states of the packaging process when only part of the genome has been pumped in. We propose that the dsDNA would adopt a figure-eight shape (“8”) at the early stage of the packaging, which can be regarded as a “relaxed” form of the rod-and-coil fold (Fig. 5a). Indeed, we have found a few intermediate-state particles in the T4 dataset that seem to support this notion. These particles have loose DNA in their capsid and with no tail. As tailed bacteriophages are known to attach their tail to their head only when the DNA packaging is complete (i.e., “headfull”), these particles are likely at an intermediate stage of genome packaging. In contrast, some other particles with almost no DNA in their capsid but with a tail are likely broken ones with their genome leaked out or ones that have been triggered for genome ejection during the purification (Fig. S6). As shown in Fig. 5, sliced views of the DNA genome in these intermediate-state T4 particles looked very similar to those of a 3D model we generated by relaxing the rod-and-coil model of the mature T4, in which the figure-eight shape is clearly visible from the direction of the portal. It was observed previously that after partial ejection of the genome, the remaining DNA in phage T4 occupies the entire volume of the capsid in the presence of monovalent cations but condenses into a compact toroid when exposed to the tetravalent cation spermine^28,29^. Our model of the intermediate state is consistent with this observation. Moreover, it could explain the easy conversion of the DNA between these two seemingly completely different states, as the figure-eight shape is topologically equivalent to the toroid.

**Fig. 5.**
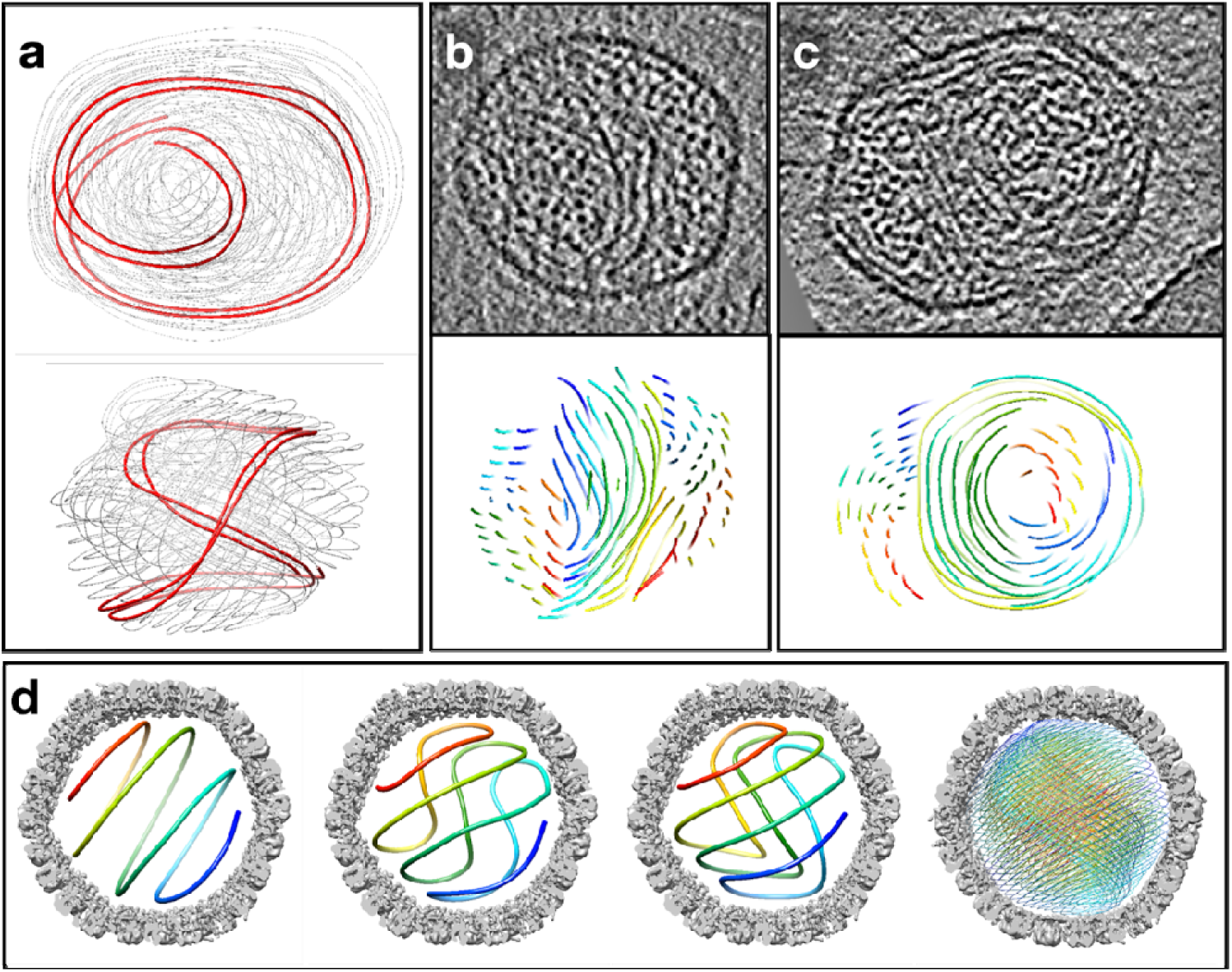
Genome packaging process in T4 and HSV-1. (a) Top and side view of the early stage DNA packaging model of T4, with a segment highlighted to show the figure-8 shape formed by dsDNA. (b-c) Comparison between the cross-section of the model in a, and the slice view of DNA inside T4 bacteriophage at intermediate packaging states. (d) Diagram of the proposed genome packaging process in HSV-1.

Extrapolating the intermediate state to even earlier time points, it is reasonable to hypothesize that the very first bit of the packaged dsDNA would have a toroidal fold. As more dsDNA is pumped into the capsid, the toroid starts to twist inward and forms the shape of figure-eight. As the figure-eight shape gets squeezed by more and more incoming dsDNA, the twists would get pushed towards the two ends, generating a straight segment in the middle to bundle and form the central rod, finally reaching the rod-and-coil form we observe in mature capsids (Fig. 5d). Obviously, in a capsid with a relatively large diameter (spherical capsid) or long axis (prolate capsid) and/or with more densely packed dsDNA, the central rod would become more prominent compared to those in a smaller capsid and/or less-densely packed genome. A most prominent example for the effect of the capsid size could be found in a “giant T4” mutant with a much-elongated capsid, where the rod-and-coil shape of the genome could be appreciated from negative staining EM micrographs of partially ruptured viral particles (see Figure 5d of reference ^5^).

This work is made possible by direct 3D visualization of individual viral particles with cryoET. The recent advances of the subtilt refinement methods greatly improve the achievable resolution of subtomogram averaging, and the information acquired from the refinement can be applied back to the raw sub-tilt series to boost the resolution of individual subtomograms. Our work demonstrate that this technique now provides an opportunity to visualize heterogeneous features and rare events in the biological systems at true molecular level.

## Acknowledgement

This work made use of instruments in the CryoEM Core Facility at Case Western Reserve University (CWRU), and computational resources in the Core Facility for Advanced Research Computing at CWRU and SLAC Shared Scientific Data Facility (SDF) at Stanford University. We acknowledge general funding support from CWRU (start-up package to X.D.) and the National Institutes of Health (R21MH125285 and R01GM150905 to M.C., R35GM151043 to X.D.).

## Author Contributions

X.D. conceived the project; X.D., X.S., B.S., Z.M., and T.T. prepared samples; X.D. collected cryoEM data; M.C. developed methods and processed the data; M.C., X.D. and Z.M. interpreted the data; M.C. and X.D. wrote the manuscript; all authors reviewed and approved the manuscript.

## Methods and Materials

### Sample preparation

Culture and purification of HSV-1 followed the same method as described before (cite Dai 2018 Science). Briefly, HSV-1 KOS strain (ATCC VR-1493) was cultured in Vero cells (ATCC CCL-81) using Dulbecco’s modified Eagle medium (DMEM) supplemented with 10% fetal bovine serum (FBS). Virion particles were harvested from the culture media, concentrated by ultracentrifugation pelleting, purified with sucrose density gradient, and resuspended in TN buffer (20mM Tris-HCl pH7.4, 150mM NaCl) for cryoEM sample preparation.

Bacteriophage T4 (ATCC 11303-B4), T7 (BAA-1025-B2) and their recommended host strains of Escherichia coli (ATCC 11303 for T4, and ATCC BAA-1025 for T7) were all purchased from ATCC and propagated following the standard protocol described on the ATCC website. Briefly, single clones of the bacteria were propagated in soft agar and inoculated with the phage stocks to make fresh, high-infectivity stocks, which were then added to log-phase bacterial broth cultures for large scale phage production. A final concentration of 10mM CaCl_2_ and 10mM MgCl_2_ were added to the culture to facilitate bacterial lysis by the phage. The lysed culture was clarified by centrifugation at 10,000g for 20min to remove the bacterial debris (repeated once) and then centrifuged at 100,000g for 2hr to pellet the phage particles. The pellet was soaked in TN buffer overnight on ice before resuspension. A continuous CsCl density gradient was prepared in parallel by centrifuging 45.41%(wt/wt) (ρ=1.5g/mL) CsCl solution (in TN buffer) overnight at 250,000g. The resuspended phage particles were then added on top of the density gradient and centrifuged at 250,000g for 2hr. The band of the phage particles was collected and dialysed overnight in TN buffer using a dialysis cassette to remove the CsCl. The phage solution was then concentrated using an ultrafiltration unit and ready for cryoEM sample preparation.

CryoEM sample freezing was done with a Vitrobot Mark IV using 200-mesh Quantifoil Cu R3.5/1 grids.

### Tomography data collection

Tomographic tilt series were collected using serialEM and the FastTomo script^30^ on a Titan Krios microscope equipped with Gatan BioQuantum K3 imaging filter and camera. A bidirectional scheme was used for the tilting, from −30° to +60° and then to −60° with a 2° step. A fixed electron dosage of about 2.5e^-^/^2^ (measured at empty area) was used for each exposure, which was fractionated into 12 frames. A nominal magnification of 81,000x was used, corresponding to a pixel size of 0.539 LJ/pixel on the camera at super-resolution mode. A target defocus of −1.6 micron was set at the beginning of each tilt series. A slit of 20 eV was used for the imaging filter.

### Tomography data processing

Movie stacks of the tilted exposures were drift-corrected using Motioncorr2^31^ and binned 2x in the Fourier space to generate micrographs with a pixel size of 1.078 LJ/pixel. A 4x3 tile with 20% overlap was used. Dosage weighting was disabled.

For the HSV-1 dataset, tilt series alignment and tomogram reconstruction were done with an automated pipeline in EMAN2. With the geometry of the tilt determined from the tilt series alignment, a CTF is estimated for each micrograph. Locations of the viral capsid particles were selected manually from the tomograms, and raw subtomograms were generated from the tilt series after per-particle-per-tilt CTF correction. From 17 tomograms of HSV-1, 41 particles were selected.

A previously published asymmetric reconstruction of HSV-1 capsid from single-particle CryoEM was downloaded from EMDB (EMD-9864) and used as an initial model for subtomogram averaging of HSV-1. The structure was low-pass filtered and phase randomized to 50Å before serving as reference for the “gold-standard” refinement. In sum, 3 iterations of subtilt translational refinement and 1 iteration of subtilt rotational refinement, combined with 4 iterations of subtomogram averaging, were performed in EMAN2. Icosahedral symmetry was applied throughout this process. After refinement, the structure of HSV-1 capsid reached 14.7 Å resolution.

Processing of the bacteriophage T4 dataset followed the same workflow as described above for the HSV-1. From the dataset, 184 particles were selected from 23 tomograms. The head part of an existing T4 structure (EMD-2774) was low-pass filtered to 50Å and used as the initial model of the refinement. The refinement of the T4 head was performed with d5 symmetry, and the structure reached 13 Å resolution. After symmetrical refinement, iterative symmetry relaxation was conducted to determine the position of the tail in each particle. After asymmetrical refinement of the T4 head, we determined the structure of the bacteriophage tail as three segments to compensate for the flexibility of tail bending. For each segment, subtomogram particles located at the lower end of the previous segment were re-extracted from the tilt series, and an averaged structure was determined from those particles with c6 symmetry. Finally, the asymmetrical structure of the virus head, as well as the three segments of the virus tail, were fused together to form the full structure of the T4 bacteriophage. The resolution of the structure ranges from 10 to 30 Å throughout the structure, and the density map was filtered according to the local resolution (Fig. S7).

For the bacteriophage T7 dataset, 1040 particles from 37 tomograms were used for the subtomogram and subtilt refinement. An existing T7 structure (EMD-1164) was low-pass filtered and phase randomized to 50Å before serving as a reference for the refinement. The subtomogram average with icosahedral symmetry reached 5.6Å (Fig. S8), with secondary structure elements matching well with an atomic model of the T7 head (PDB 3J7X). Afterward, we performed two rounds of symmetry expansion and iterative classification to relax the icosahedral symmetry. The first round relaxed the icosahedral symmetry to c5 to identify the unique vertex where the portal complex and the tail are located. The second round of symmetry relaxation focused on the portal complex and relaxed the c5 symmetry to c1, resolving the symmetry mismatch between the capsid and the portal. Finally, another round of subtomogram refinement was performed on the entire particle without symmetry, using the symmetry-relaxed c1 structure and orientation as a starting point. This resulted in a subtomogram average of the entire T7 particle at 15.1 Å resolution.

In all three cases of HSV-1, T4, and T7, results from symmetrized subtilt refinement of viral capsids were used to generate polished subtomograms to better resolve the packaged dsDNA genome. Specifically, rotation and translation parameters determined from subtilt refinement were applied back to each subtilt image, and polished subtomograms were reconstructed from those 2D images Afterwards, each subtomogram was rotated back to their orientation in the original tomogram, with the zero tilt image on the x-y plane, so they can be visualized in 2D slice by slice through the z-axis. The polished subtomograms were band-pass filtered to 20-100Å to boost the contrast of dsDNA strands.

### Genome packaging parameters calculation

To measure the distance between nearest neighbor DNA duplexes, we first annotated the DNA segments that run parallel to the electron beam, because they showed up as well-separated dots in x-y plane slice views. The annotation was performed on polished subtomograms using the convolutional neural network based semi-automated tomogram annotation protocol provided in EMAN2. Coordinates of the DNA cross-sections were selected from the annotation, and further refined by locally fitting a Gaussian to the 2D slice view of the DNA. For each point of the DNA cross-section, we measured the distances to its nearest DNA duplexes on the same x-y plane. Finally, the distribution of these nearest neighbor measurements was fitted with a Gaussian to estimate the average distance between DNA duplex pairs. For the HSV-1 particles, 5097 DNA cross-sections were measured, and the average nearest neighbor distance was 31.3 Å, with a standard deviation of 3.7 Å. For particles of fully assembled T4 bacteriophage, 2066 DNA cross-sections were measured, and the distance was 25.6±4.0 Å. For particles of T4 assembly intermediates, the sample size was 2145, and the measured distance was 40.3±8.0 Å. For the particles of T7 bacteriophage, 1177 DNA cross-sections were measured, and the distance was 23.8±2.9 Å.

From the measured local distance of DNA duplexes, we calculated the total genome length as a validation of our measurement. The total volume inside the capsid is estimated to be 4.49e8 Å^3, measured from the averaged structure of the capsid. Calculating from the local arrangement, a DNA hexagon pillar with a height of 34Å would have a cross-section area of 2540 Å^2, and a volume of 8.64e4 Å^3. For B-form DNA, the hexagon pillar would contain 3 unique 10bp strands, with a total of 30bp. Combined with the total volume inside the capsid, the total length of dsDNA inside the capsid is estimated to be 156kbp, close to the actual HSV DNA length of 152kbp.

Finally, we estimate the number of turns the coil DNA makes per each DNA strand on the rod. From the particles whose central rod is perpendicular to the imaging plane, we measure the area of the central rod to be around 74000 Å^2. Assuming the rod extends throughout the capsid, the volume of the DNA rod is 6.73e7Å^3. So the volume ratio between DNA on the coil and DNA on the rod is 5.68:1. As the local packaging of DNA strands is as tight on the coil as it is on the rod, this should also be the ratio of the length of DNA on the coil and that on the rod. Here, we use a simplified model of DNA in a spherical capsid, the rod DNA raises straight along z, whereas the coil DNA wraps around following the spherical shell. In this model, we can compute the length ratio between the coil and rod DNA as a function of the number of turns the coil DNA makes around the rod. For example, the coil-rod length ratio would be 3.13 if the coil makes one turn (360 degrees) around the rod. From the model, when the coil makes 2.11 (760 degrees) turn per each rod, the volume ratio matches the experimental estimation. Here we round it down to 2 for simplification when generating the HSV DNA organization model.

### DNA model building

From the packaging parameters estimated from the subtomograms, we build a conceptual model of DNA organization of HSV. We orient the DNA rod along the z axis, and determine the number of DNA strands per each layer of the coil by the volume of each coil layer. Then build the coil DNA starting from the outermost shell. An 1D exponential function is used to represent the path of a DNA coil along the z direction, so the rise is small near zero, and sharp near one, where the seam line locates. A number of those curves are put together with even spacing along z, corresponding to the number of strands in the rod for that layer. Then, we warp the 1D curves along a cylinder for two cycles, forming the initial shape of the coil DNA. Afterward, the two ends of the DNA coils are connected to the corresponding points at the ends of the central rod, and the model is mapped to the surface of a sphere. From this model, we run a local optimization, so the DNA fits in the inner surface of the capsid, and the DNA strands have an even spacing locally. After building the model for the outermost layer, we repeat the process for each of the inner layers. The operation for each layer is largely the same, except for the later layers, instead of fitting the DNA to the inner surface of the capsid, the DNA is fit to the inner surface of the previous coil layers. Finally, after building the model of all layers, the DNA is connected at the central rod to form a single strand, and local conflicts between layers are resolved manually in UCSF Chimera (Fig S5).

To model the packaging process, we simplify the DNA organization model to a single closed loop. We use a closed circle as the start point of packaging, and select one strand in the central rod from the complete DNA organization model, as well as its corresponding coil, as the end point of the packaging process. For the end point model, the two ends of the path are connected at the central rod, so it also forms a closed loop. Then, we search for the best relative orientation between the start and end point, so the morphing path between the two models is minimized. From this orientation, we can generate a movie of the morphing process from a closed circle to the rod-and-coil fold of the final packaging model (Supplementary video 12). Finally, for each frame of the movie, we populate a full DNA model around the closed loop according to the local DNA arrangement pattern.

To build the DNA model of the T4 assembly intermediate states, we start from the one model from the DNA packaging movie built for HSV. The model is first orientated so that the central rod of the model is aligned with the rod in T4 intermediate state particles. Then, we morph the model so the shape of the envelope of the DNA model matches the shape inside the capsid of T4. Finally, the model is adjusted locally in Chimera to resolve some conflicts between strands, and the local orientation of the curves is aligned with the density of DNA in individual particles.

---

**Fig. S1.**
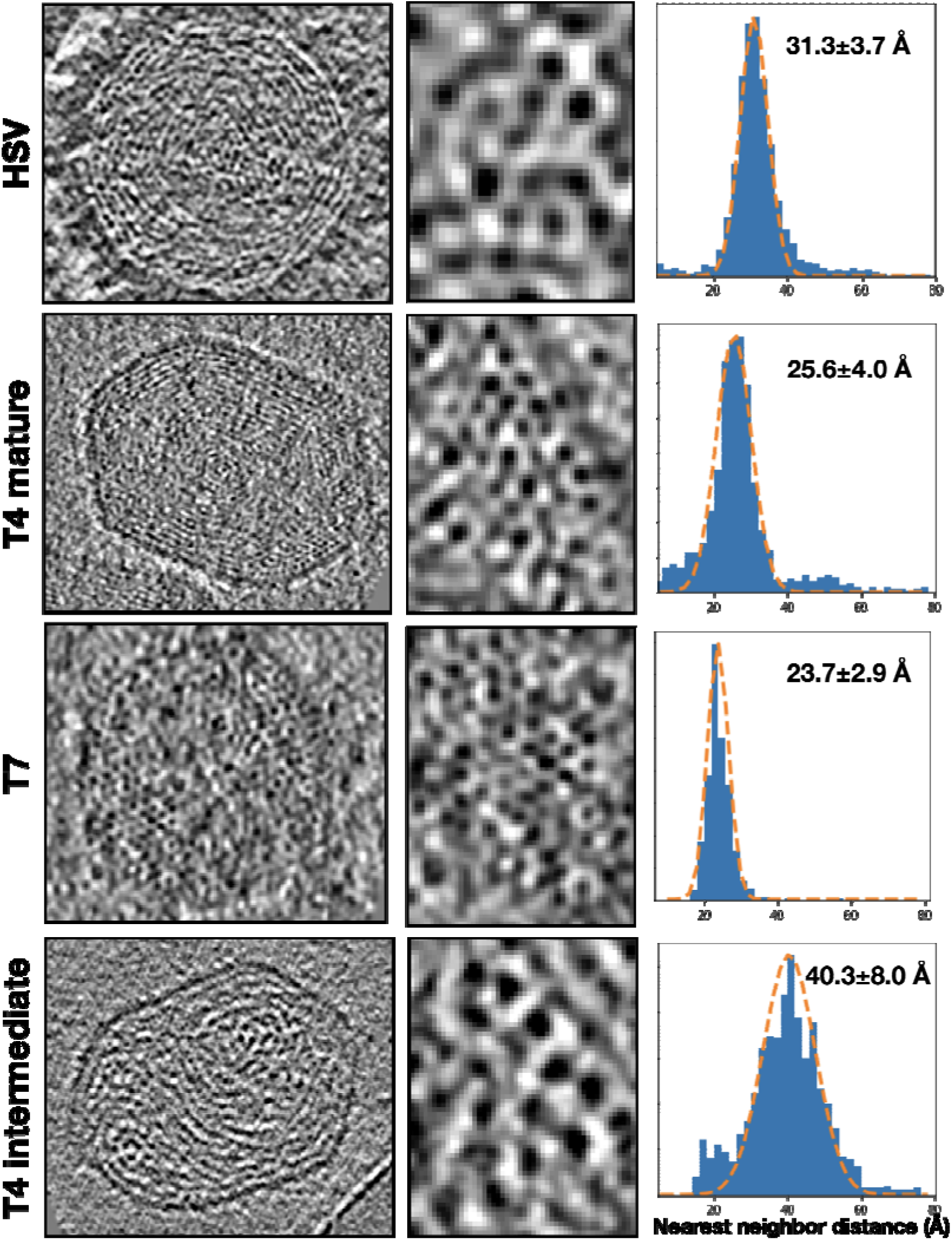
Distribution of the distance between the nearest dsDNA strains in different viruses and different assembly states. Left column: slice view of polished subtomograms. Middle column: zoomed-in view of DNA bundle. Right column: Histogram of nearest neighbor distance distribution, with fitted Gaussian distribution overlaid.

**Fig. S2.**
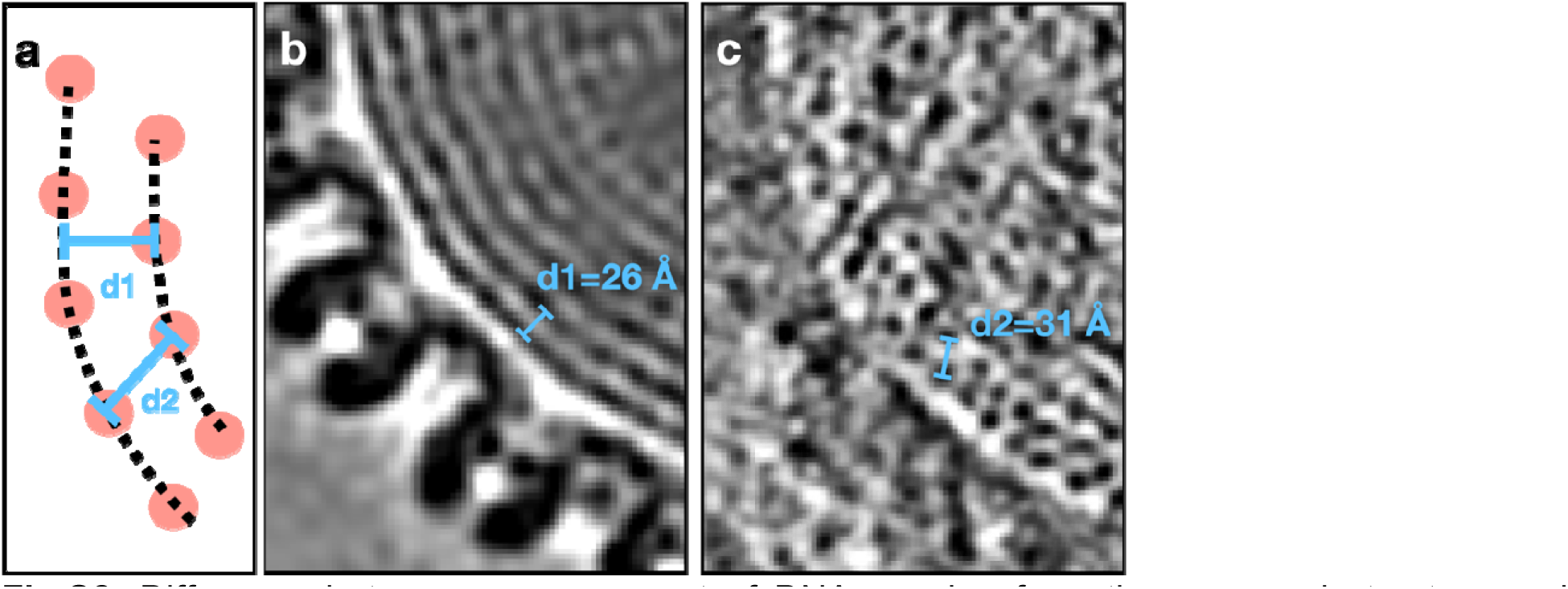
Difference between measurement of DNA spacing from the averaged structure and individual particles. (a) Diagram showing the difference between the two measurements. (b) Measurement of DNA spacing from the averaged structure of HSV-1. (c) Measurement of DNA spacing from one polished subtomogram of HSV-1.

**Fig. S3.**
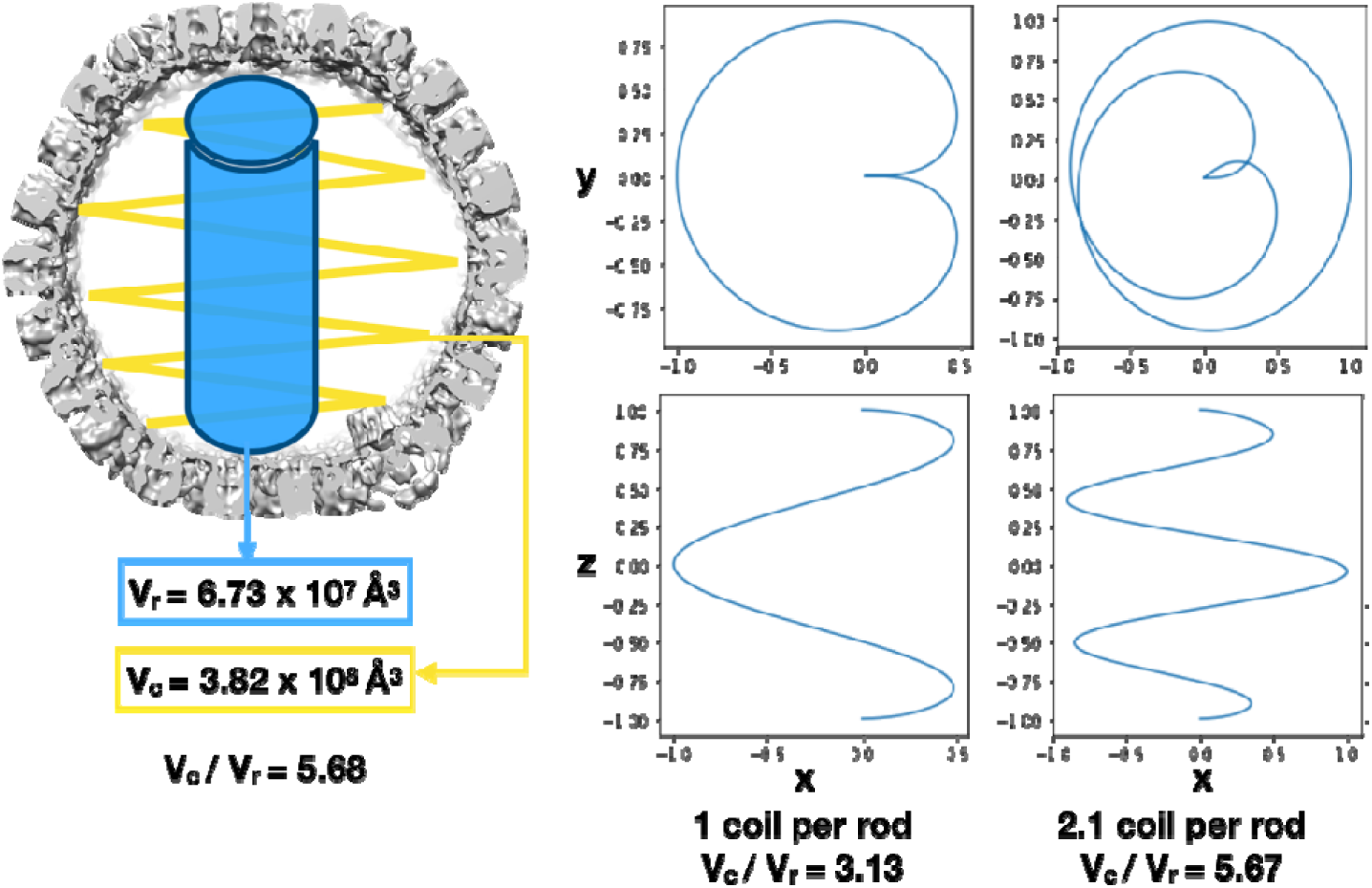
Calculation of the length ratio between dsDNA in the coil versus the rod fold in HSV. The number of coils per rod is determined so that the volume ratio of DNA within the rod and coil matches the observation from HSV particles.

**Fig. S4.**
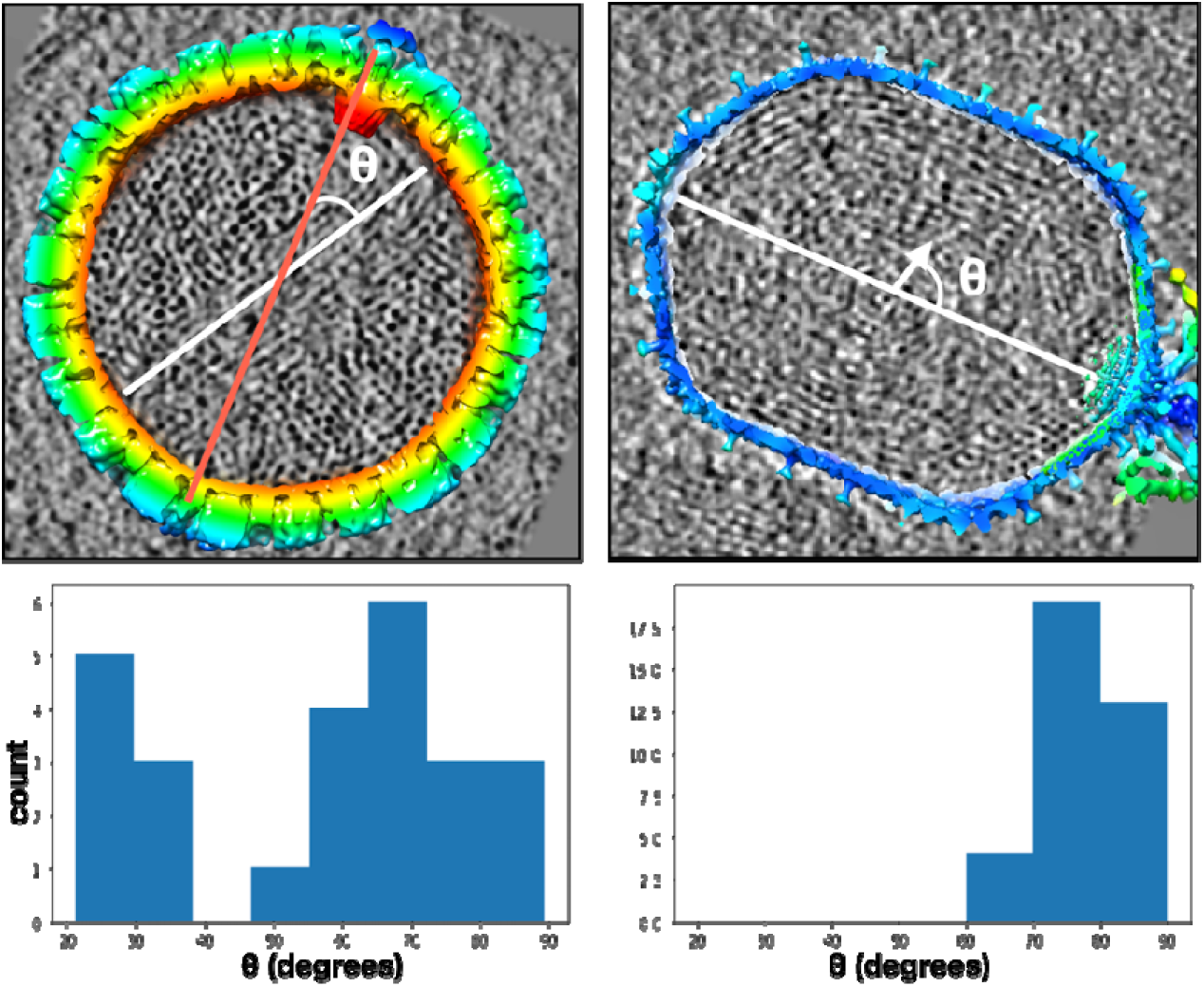
Illustration of the angle between central DNA rod and portal axis in HSV-1 (left) and the T4 bacteriophage (right), and the distribution of the angle among the particles.

**Fig. S5.**
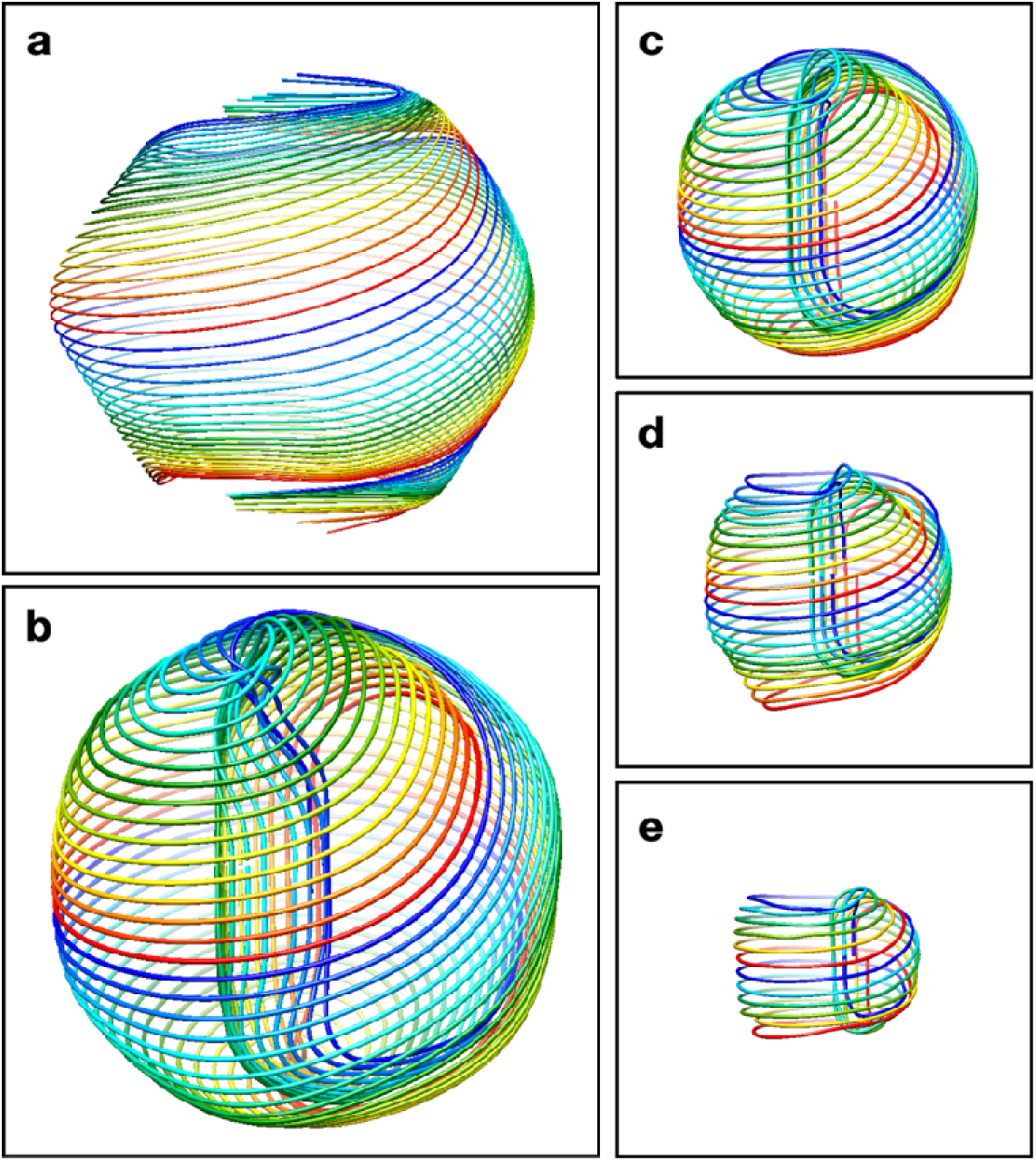
Generation of dsDNA packaging model for HSV-1. (a) Initial path of the DNA strands of the outermost coil layer fit to the shape of the capsid. (b) Final path of the outermost coil layer and corresponding strands in the central rod. (c-e). Final paths of the 3,5,7th coil layers and corresponding strands in the central rod.

**Fig. S6.**
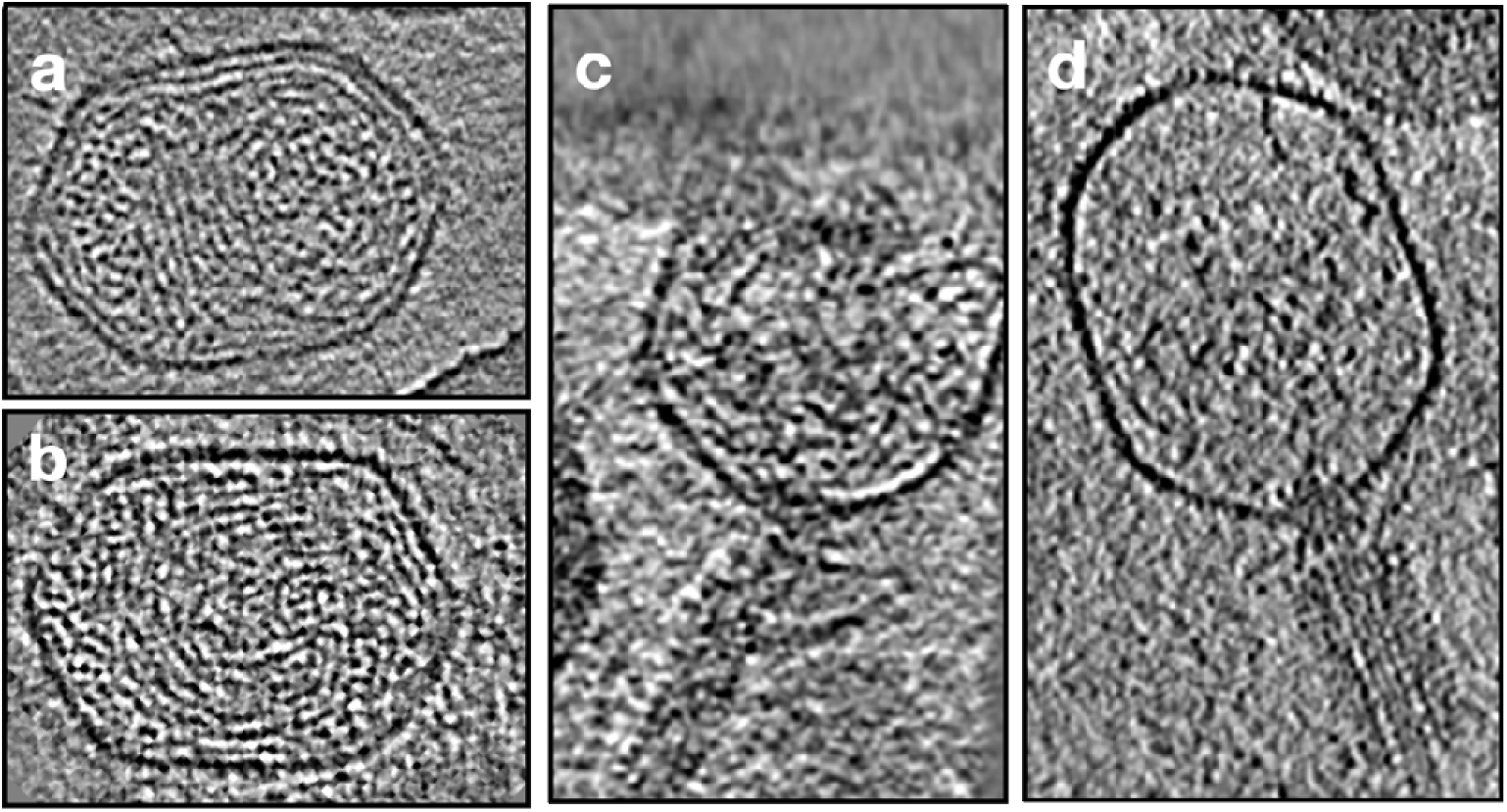
Comparison of T4 genome packaging/releasing intermediates. (a-b) Slice view of T4 particles during the DNA packaging process. The genome density inside the capsid is lower than that of the mature bacteriophage, but the organization is ordered. (c-d) Slice view of T4 particles during the DNA releasing process, showing the disordered DNA inside.

**Fig. S7.**
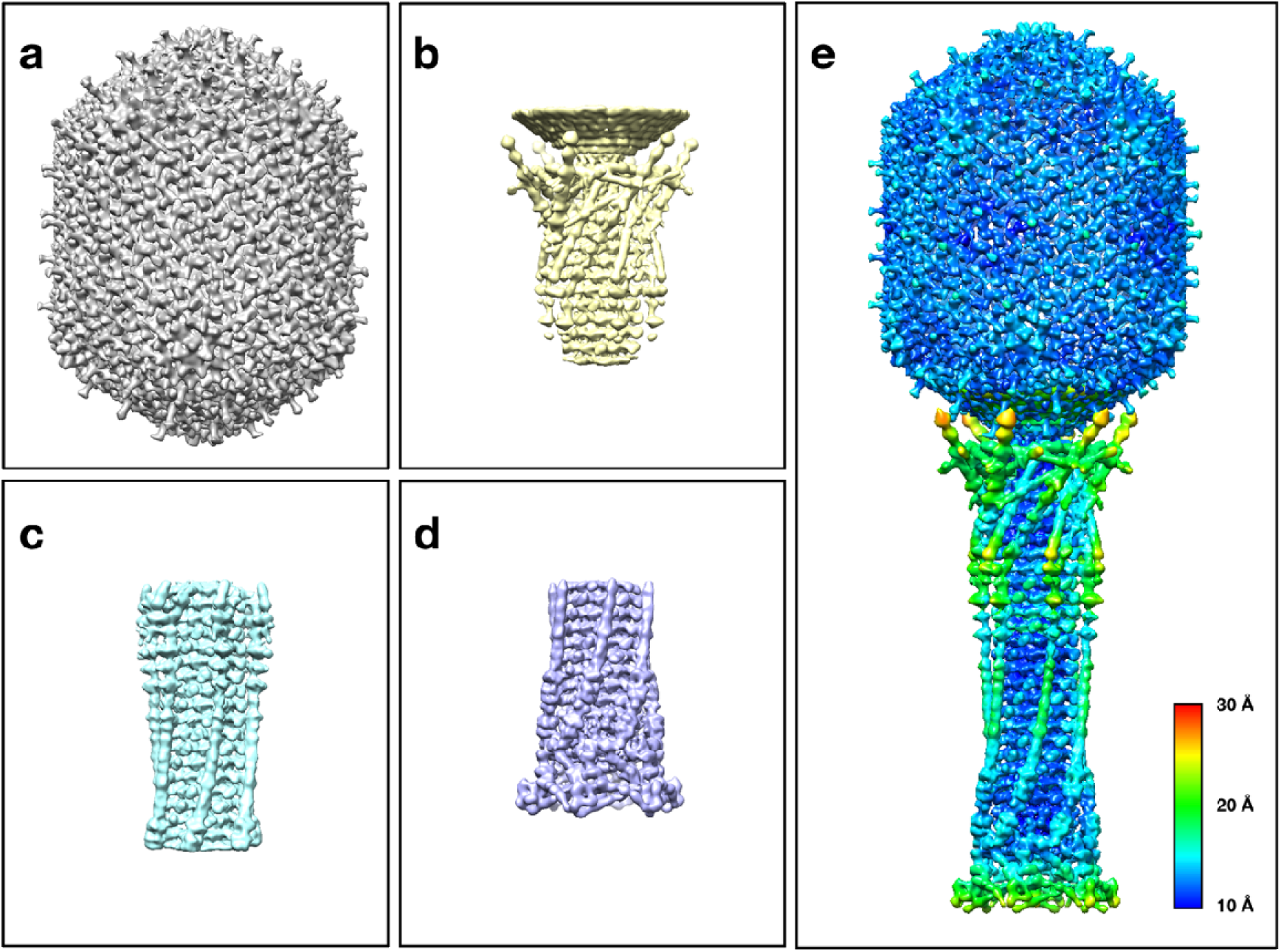
Structure of T4 bacteriophage. (a-d) Subtomogram averages of virus capsid and three segments of the virus tail. (e) Merged structure of the entire bacteriophage, colored by local resolution.

**Fig. S8.**
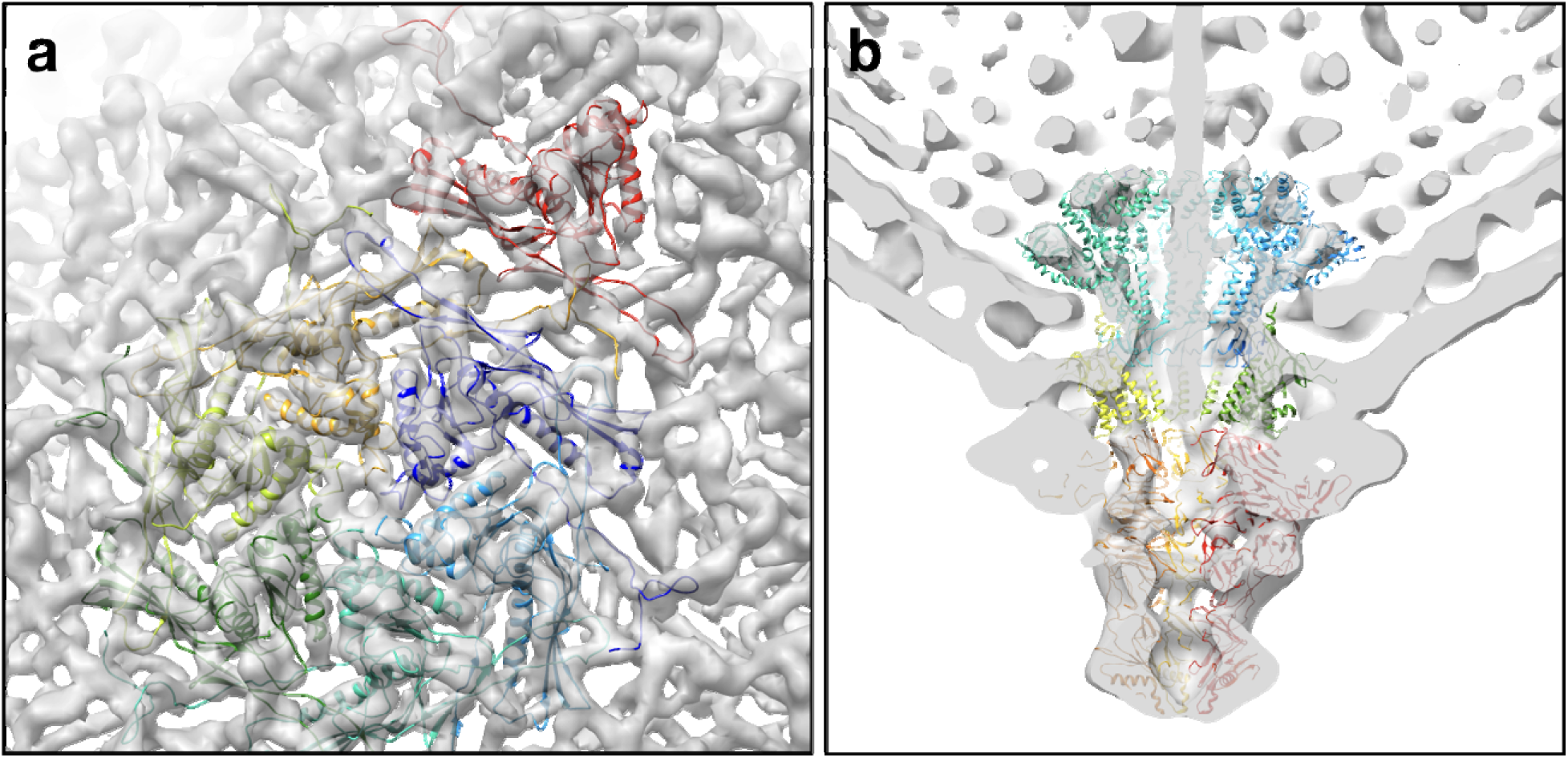
Structure of T7 bacteriophage capsid. (a) Capsid structure determined with icosahedral symmetry with corresponding high resolution model (PDB: 3j7x) fitted into the density. (b) Capsid structure determined without symmetry, showing the portal complex. The corresponding high resolution model (PDB: 6r21) fitted into the density.

Supplementary video 1 - Tomogram slice view of HSV-1

Supplementary video 2 - Slice view of polished HSV-1 particle #1

Supplementary video 3 - Slice view of polished HSV-1 particle #2

Supplementary video 4 - Tomogram slice view of T4 bacteriophage

Supplementary video 5 - Slice view of polished mature T4 particle #1

Supplementary video 6 - Slice view of polished mature T4 particle #2

Supplementary video 7 - Slice view of polished intermediate T4 particle #1

Supplementary video 8 - Slice view of polished intermediate T4 particle #2

Supplementary video 9 - Slice view of polished T7 particle #1

Supplementary video 10 - Slice view of polished T7 particle #2

Supplementary video 11 - 3D view of the DNA packaging model in HSV-1

Supplementary video 12 - 3D view of the morphing from a DNA toroid to the rod-and-coil fold

